# Genome-Wide Classification of Type I, Type II and Type III Interferon-Stimulated Genes in Chicken Fibroblasts

**DOI:** 10.1101/793448

**Authors:** Diwakar Santhakumar, Mohammed A. Rohaim, Muhammad Munir

**Affiliations:** Division of Biomedical and Life Sciences, Faculty of Health and Medicine, Lancaster University, Lancaster, LA1 4YG, UK

**Keywords:** interferons, chicken, birds: innate immunity, ISGs, IFN regulated genes, immunity

## Abstract

Interferons (IFNs) play central roles in establishing innate immunity and mediating adaptive immunity against multiple pathogens. Three known types of IFNs identify their cognate receptors, initiate cascades of signalling events and eventually result in the induction of myriad of IFN-stimulated genes (ISGs). These ISGs perform multitude of functions and cumulatively corroborate a bespoke antiviral state to safeguard hosts against invading viruses. Owing to unique nature of chicken’s immune system and lack of foundational profiling information on the nature and dynamic expression of IFN-specific ISGs at the genome scale, we performed a systematic and extensive analysis of type I, II and III IFN-induced genes in chicken. Employing pan-IFN responsive chicken fibroblasts coupled with transcriptomics we observed an overwhelming over-representation of up-regulated ISGs by all types of IFNs. Intriguingly, prediction of IFN-stimulated response element (ISRE) and gamma-IFN activation sequence (GAS) revealed a substantial number of GAS motifs in selective significantly induced ISGs in chicken. Extensive comparative, genome-wide and differential expression analysis of ISGs under equivalent signalling input catalogue a set of genes that were either IFN-specific or independent of types of IFNs used to prime fibroblasts. These comprehensive datasets, first of their kinds in chicken, will establish foundations to elucidate the mechanisms of actions and breadth of antiviral action of ISGs which may propose alternative avenues for the targeted antiviral therapy against viruses of poultry and public health importance.

## 1. Introduction

Innate immunity defines the first line of host defense, and interferons (IFNs) play central roles in mediating host resistance against viruses, bacteria, fungi and parasites by directly and indirectly regulating multiple immune cells [1]. IFNs are part of the class II cytokine family and are divided into three types (I, II and III) based on their receptor specificity, sequence homology, evolutionary relatedness, functional activity of these cytokines and nature of interferon-stimulated genes (ISGs) induction [2,3]. The ISGs display a spectrum of cellular functions and the cumulative actions of these ISGs define the antiviral state of the hosts.

Amongst all IFNs, type I IFNs were first identified, most studied, are diverse and potent antiviral cytokines. Compared to multiple IFNs in human, only two well-characterized type I IFNs (IFN-α and IFN-β) are reported in chicken and we have recently identified a third novel chicken type I IFN (IFN-κ) that carried antiviral activities both *in vitro* and *in ovo* [4]. While all type I IFNs are structurally diverse, they bind to the same heterodimeric receptor complex composed of one chain of the IFN-α receptor 1 (IFNAR1) and one chain of the IFN-α receptor 2 (IFNAR2). In contrast, type II IFN, constituted of single gene encoding IFN-γ, interacts with receptors complexes of IFNγR1 and IFNγR2 and play pleiotropic roles in regulating the maturation and differentiation process of several immune cells and activating T helper 1-type immune responses [5]. More recently identified class of IFNs are type III IFNs or IL-28/29 which are encoded by a single gene of IFN-λ in chicken compared to four genes in human [4]. The type III IFN interacts with heterodimeric receptor complex of IL-28Rα and IL-10Rβ, which are predominantly expressed on the epithelial cells of the host.

Upon binding to their cognate receptors, all types of IFNs initiate a cascade of intracellular events that culminate in the induction of several hundreds genes; however each type of IFN carries a certain level of distinction [4]. In spite of type I and III IFNs are sharing many properties, recent discoveries have revealed context-specific functional differences [6]. It has been shown that type I and III IFNs in mammals initiate the same signalling pathway through phosphorylation of Signal transducer and activator of transcription 1 (STAT1) and STAT2 heterodimers possibly by tyrosine kinase 2 (TYK2) and Janus Kinase 1 (JAK1). However, type II IFN triggers this pathway via the activation of STAT1 homodimers mediated by JAK1 and JAK2 kinases [4].

Mammalian type I and type III IFNs signal through the JAK-STAT pathway to activate the heterotrimeric interferon-stimulated gene factor 3 (ISGF3) complex, comprised of phosphorylated STAT1 and STAT2, and interferon regulatory factor 9 (IRF9) [7,8] (yet to be identified in chickens). In contrast, type II IFNs initiate the formation of a STAT1–STAT1 homodimer to assemble GAF, without the need of IRF9. Upon nuclear translocation, ISGF3 and gamma-interferon activation factor (GAF) bind to IFN-stimulated response elements (ISREs) [6] or gamma-IFN activation sequence (GAS) motifs, respectively^7^ leading to the transcriptional activation of hundreds of ISGs which encode proteins that act via a variety of mechanisms to restrict viral infection [9-11].

It has been demonstrated that JAK-STAT pathway initiates transcriptional regulation of a string of cytokines, chemokine, antimicrobial products and regulators of apoptotic pathways [2,7]. In mammals, a range of genome scale transcriptomics screening approaches has been applied to map the diversity, kinetics and expression patterns of these antiviral factors. Based on these data, recent studies have applied large-scale screening platforms to elucidate functional importance of these ISGs regulated by IFN-α, IFN-λ and IFN-γ [10,11]. However, primary focus of these studies was restricted to type I and type II IFNs whereas growing evidences are highlighting the crucial roles of IFN-λ (type III IFN) in establishing antiviral state. Recently, it has been demonstrated that paracrine singling of IFN-λ is fundamental in early innate immune responses and to inhibit influenza virus spread [12], which pose enormous zoonotic (e.g. H7N9, H5N1, H9N2 and H5N2) threats to public health and remains the most economically devastating disease in the poultry [13].

Most of our understandings on ISGs-mediated antiviral functions in chicken are driven from human and other mammalian studies. Given the fact that chicken differs fundamentally from other mammals in innate immunity, majority of current concepts are mainly based on assumptions. Some out of many examples include absence of RIG-I, IRF9 and IRF3 homologues, lack of anti-viral roles of chicken Mx, absence of many essential molecules in the innate immune signalling pathways such as IFIT1/2/3, shorter versions of ZC3HAV1 and many of the DNA sensors [2,3]. Intriguingly, despite the absence of these key components, chickens respond to highly pathogenic avian influenza (HPAIVs) and other viruses, and mount potent type I IFN responses [14-16] highlighting the presence of yet unknown and evolutionary compensatory mechanisms in chicken.

Given the lack of comprehensive data on the nature of type I, II and type III IFN-mediated antiviral profiling, uniqueness of the chicken as model system, and being an important poultry species, studies have started to catalogue IFN-induced and IFN-regulated genes in chicken [17,18]. However, these studies have exclusively identified expression dynamics of ISGs induced by type I IFNs and nature and regulation of type II and type III mediated ISGs are required to be explored in chicken at genome-wide scale. Additionally, a side-by-side comparison of the type I, II, and III IFN-driven host cell transcriptomes has not been reported so far in chicken. Therefore, in order to lay foundation for future research on functional annotation of ISGs, we performed intensive gene expression analysis in pan-IFN responsive chicken fibroblasts and provide a comprehensive and comparative data on the chicken ISGs stimulated with type I, II and III IFNs. This data will help to facilitate underpinning the comparative studies with human ISGs and to stimulate large-scale functional screens as has been performed in mammals [10].

## 2. Materials and Methods

### 2.1. Cells culture and media

Immortalized chicken fibroblasts (DF-1) and human embryonic kidney cells 293T (HEK-293T) cells were maintained in Dulbecco’s modified eagle medium (DMEM) (Gibco, Carlsbad, CA, USA) supplemented with 10% foetal bovine serum (FBS), 1% penicillin and streptomycin (P/S) at 37 °C in 5% CO2 incubator.

### 2.2. Production of Interferons

The open reading frame (ORF) sequences for chIFN-α, chIFN-γ and chIFN-λ were synthesized (GeneArt, Thermo, UK) and cloned into the pcDNA3.1+ vector. The pcDNA3.1+ encoding chIFN-α, chIFN-γ and-chIFN-λ plasmids were transfected into HEK-293T cells using the Lipofectamine 2000 Transfection Reagent (Invitrogen). Supernatants were collected at 48 h and 72 h post-transfection, then were cleared by centrifugation (1500 rpm for 5 min) and pooled and quantified using IFN-bioassay.

### 2.3. IFN bioassay

IFN-induced protection against VSV-GFP was used to identify IFN-producing stable clones (chIL28RA) and to quantify IFN preparations, as described before [19] and we reported earlier [3]. Briefly, DF-1 cells were seeded in 96-well plates until they are 90% confluent and treated with serial dilutions of supernatants containing interferons for 24 hours. These interferon-stimulated cells were infected with VSV-GFP (MOI of 0.1). After 24 hours post-infection, VSV-GFP replication was correlated with the change in GFP fluorescence signal intensities using Luminometer (Promega, Madison, WI, USA). The percentage antiviral activities of IFNs were determined by comparing the percentage reduction of corrected GFP signal intensity (GFP signal intensity of IFN treated and virus infected wells minus background fluorescence signal intensity of uninfected control) with the mock-treated and VSV-GFP-infected control wells. One unit (U) of IFN in the tested IFN preparations was defined as the volume containing 50% inhibitory activity against VSV-GFP. A total of 1000 UI of IFNs were used for stimulation of chIL28RA DF-1 cells.

### 2.4. Library preparation for mRNA sequencing

Total RNA was extracted from IFN-primed or mock treated chIL28RA DF-1 cells in duplicate for RNA-sequencing using the TRIzol reagent (Invitrogen, USA). Approximately 2.5 µg of RNA per sample was used as input material while the mRNA was enriched by Oligo (dT) beads and then split into short fragments using fragmentation buffer (Thermo) that were then reverse transcribed into cDNA using random primers. Second-strand cDNA was synthesized by DNA polymerase I, RNase H, and dNTP. The cDNA fragments were then purified using the QiaQuick PCR extraction kit, poly(A) tails were added, and the ends were repaired and ligated to Illumina sequencing adapters. The ligation products were size selected by agarose gel electrophoresis, PCR amplified, and were then sequenced using the Illumina HiSeq in 2×150bp configuration by Genewiz Co. (New York, USA).

### 2.5. RNA-Seq data analysis

Sequence reads were trimmed to remove possible adapter sequences and nucleotides with poor quality using Trimmomatic v.0.36 [20]. The reads were then mapped to the chicken reference genome available on ENSEMBL using the STAR aligner [21]. The RNA-seq aligner was executed using a splice aligner, which detects splice junctions and incorporating them to help align the entire read sequences which generate BAM files. Unique exon hit counts were calculated using feature counts from the Subread package. After mapping and total gene hit counts calculation, the total gene hit counts table was used for downstream differential expression analysis using EdgeR [22] (http://www.rproject.org/). Expression values from each sample were carried out to estimate the sample distances. Shorter the distance, more closely related the samples are which used to identify if the two groups are closely related or not. Volcano plot analysis used to show the global transcriptional change across the groups [23].

Gene ontology analysis was conducted using GeneSCF [24]. Significant genes (absolute Log2 Fold Change, FC, >2) were first annotated with the Gene Ontology Biological Process information from the “goa_chicken” database. Then Fisher’s exact test was conducted on each ontology. A list containing statistics of over- or under-representation of GO Biological Processes relative to the unfiltered gene list was generated for each comparison and included as a deliverable. After mapping and total transcript hit counts calculation, the total transcript hit counts table was used for downstream differential expression analysis using DEXSeq [25]. To estimate the expression levels of alternatively spliced transcripts, the splice variant hit counts were extracted from the RNA-seq reads mapped to the genome. Differentially spliced genes were identified by testing for significant differences in read counts on exons (and junctions) of the genes using DEXSeq with an adjusted p value of 0.05.

### 2.6. Quantitative Reverse Transcription-PCR and Cytokine Expression

Total RNA was extracted from chIFN-α, chIFN-β and chIFN-λ (1000 UI) stimulated chIL28RA DF-1 cells using TRIzol reagents (Invitrogen, Carlsbad, CA, USA). Quantity and purity of extracted RNA was determined with a NanoDrop 1000 (PEQLAB Biotechnologie GMBH, Erlangen, Germany), while the RNA quality was analysed using a 2100 Bioanalyzer® (Agilent Technologies, Böblingen, Germany). A total of 200 ng of RNA was used in PCR reactions using SuperScript® III Platinum® SYBR® Green One-Step qRT-PCR Kit (Invitrogen, Carlsbad, CA, USA). The abundance of specific ISG mRNA was compared to the 28S rRNA in the Applied Biosystems Prism 7500 system. The reaction was carried out in ABI 7500 light cycler using the following thermo profile; 50 °C for 5minutes hold, 95 °C for 2minutes hold, followed by 40 cycles of 95 °C for 3 seconds and 60 °C for 30 seconds. Melting curve was determined at 95 °C for 15 seconds, 60 °C for 1minute, 95 °C for 15 seconds and 60 °C for 15 seconds.

### 2.7. Prediction of ISREs and GAS elements and Promotor analysis

In order to investigate if the significantly induced ISGs by the chicken type I, II, and III carry ISRE and GAS elements in their promoter regions, we scanned the 4 kb upstream promoter regions according to the approach described before [18,26,27]. Screening was performed based on Ensembl ID of most up-regulated genes for the GAS and ISRE using a Shell Script (kindly provided by Thomas Lütteke, Germany) [26].

### 2.8. Luciferase reporter assays

To determine responsiveness of chicken ISRE and GAS to chicken chIFN-α, chIFN-γ or chIFN-λ, chIL28RA DF-1 cells were grown in 96-well plate format at 2 ×10^4^ to 4×10^4^ cells/well and were co-transfected with 10 ng/well of a plasmid constitutively expressing Renilla luciferase (phRL-SV40; Promega, Madison, WI, USA) and 150 ng/well of full length pchMx or pchIFIT5 constructs. The pGL3.1 basic vector was used as a negative control. All transfections were performed using Lipofectamine 2000 and luciferase reporter assays were performed on the monolayers of chIL28RA DF-1 cells according to the manufacturer’s protocol. At 24 h post-transfection, the transfected DF1 cells were stimulated with chIFN-α, chIFN-γ, and chIFN-λ then lysed using 20µl 1× passive lysis buffer (Promega, Madison, WI, USA) at 6 h post IFN treatment, and samples were assayed for Firefly and Renilla luciferase activities using the Dual-Luciferase Reporter Assay System by following supplier’s instructions.

### 2.6. Statistical analyses

Pairwise comparisons of treated and control groups were performed using Student’s *t*-test. All statistical analysis and figures were conducted in the GraphPad Prism (GraphPad Software, La Jolla, CA, USA).

## 3. Results

### 3.1. Establishment of a Model Stable Chicken Cell Line Responsive to Type I, II and III IFNs

The aim of the present study was to reveal a complete picture of the antiviral type I, II and III-responsive genes in chicken fibroblasts. To elucidate the overall antiviral potency and the unimpeded ability of chicken IFNs to establish an antiviral system, we expressed chIFN-α, chIFN-λ and chIFN-γ in the heterologous mammalian (HEK 293T) cells as we described before for the chIFN-β and chIFN-κ [3,4]. The biological activities and responses of chicken embryo fibroblasts to these cytokines were assessed using highly sensitive and IFN-activated chMx promoter [3,26]. The chMx promoter carries *cis-acting* regulatory elements including GAS and ISRE, which are fundamental for regulation of the pan-IFN mediated transcriptional initiation.

Using chMx promoter-based reporter system, fold induction of downstream luciferase gene was measured under chIFN-α, chIFN-λ and chIFN-γ stimulation and compared to mock-treated control. The recombinant chIFN-α and chIFN-γ induced as high as 19 and 14 folds luciferase activities, respectively. Intriguingly, chIFN-λ activated a weaker chMx promoter activity in established chicken fibroblasts (Figure 1A). Since all three types of IFNs activate JAK-STAT pathway in a receptor-specific manner, it has been revealed that DF-1 cell line showed suppressed expression of chIL28Rα receptor [28], which led to attenuated promoter activities. Moreover, we demonstrated a suppressive response of DF1 against chIFN-λ in an IFN-bioassay. When wild type DF-1 cells were stimulated with chIFN-α and chIFN-γ, a substantial inhibition in the replication of an IFN-sensitive VSV-GFP virus was observed in a dose dependent manner (Figure 1B). In contrast, these cells failed to inhibit the replication kinetics of VSV-GFP, further confirming the suppressive expression of cognate receptors for chIFN-λ.

**Figure 1.**
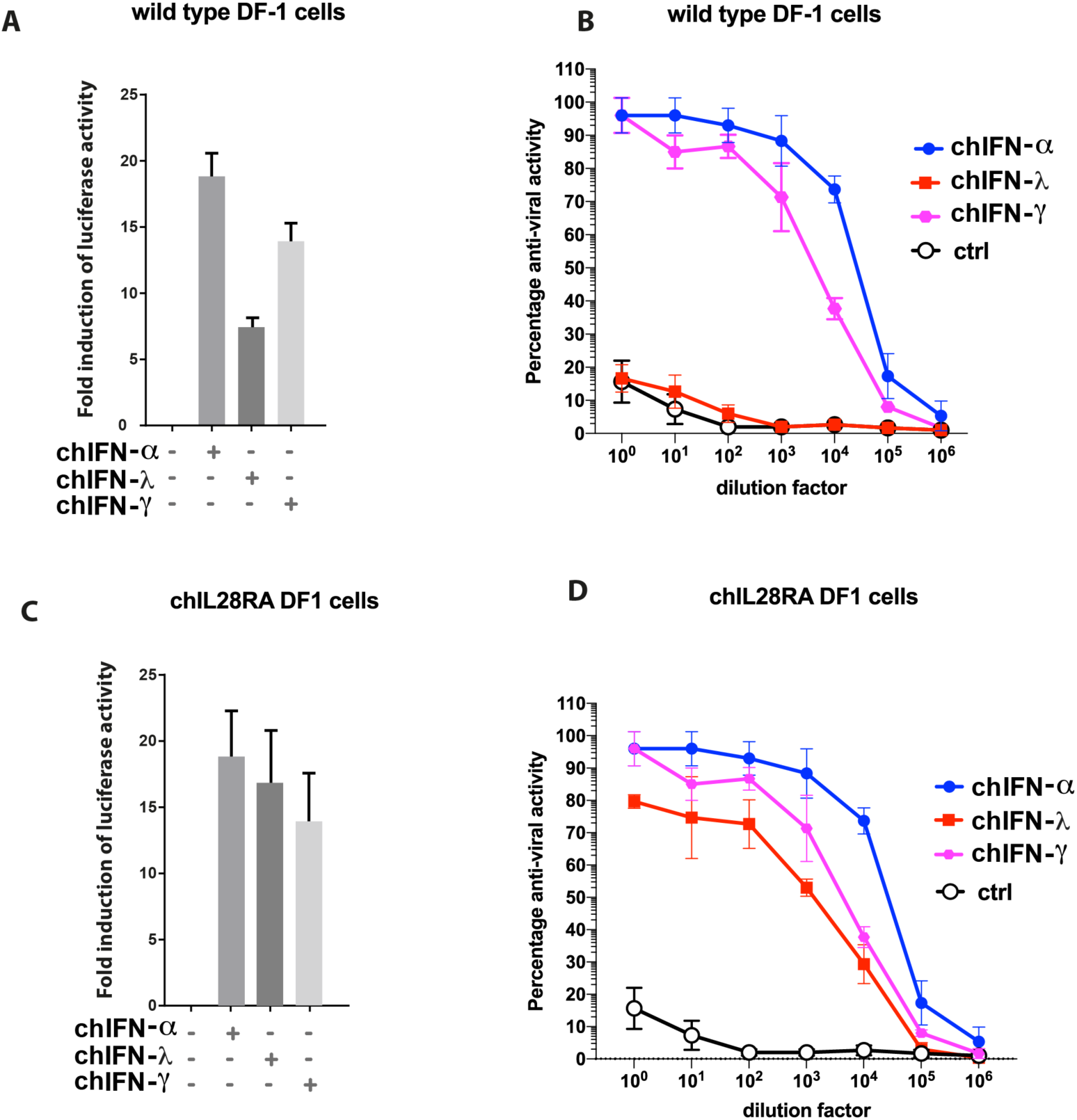
Responsiveness of chIL28RA DF-1 cells to type chicken I, II and III IFNs. Wild type (**A**) or chIL28RA DF-1 cells (**C**) were transfected with plasmid encoding luciferase genes downstream to chMx promoter and stimulated with chIFN-α, chIFN-γ or chIFN-λ for 24 hours before lysis and measuring the luciferase activities. Fold induction was calculated compared to mock-stimulated control. Wild type (**B**) or chIL28RA DF-1 cells (**D**) were treated with increasing concentration of chIFN-α, chIFN-γ or chIFN-λ for 24 hours followed by infection with the VSV-GFP for 24 hours. The level of expression of GFP was measured and plotted. The data represent three independent experiments performed in triplicate.

In order to demonstrate the supplementation of chIL28Rα can reconstitute the functional abnormalities, we used a stable DF1 cell line, which constitutively expressed chIL28Rα receptors (named as chIL28RA DF1). Stimulation of these cells with chIFN-λ profoundly activated the chMx prompter activities at the level that was comparable to chIFN-α and chIFN-γ (Figure 1C). Moreover, the chIFN-λ substantially inhibited the replication of VSV-GFP in stable chIL28RA DF1 cells. While the level of virus inhibition was lower than chIFN-α and chIFN-γ, comparative to wild type DF1, a considerable improvement was observed (Figure 1D). These results demonstrate that chIL28RA DF1 cell line is an improved system to quantify all types of IFNs, possesses a functional IFN system, and is a suitable model cell line to map the nature and dynamics of ISGs induced by all type I, II and III IFNs. Therefore, this stable cell line was used for chicken transcriptomes throughout the study.

### 3.2. Experimental Outline to Measure Chicken Transcriptomes

The recombinant chIFN-α, chIFN-λ and chIFN-γ were quantified in a VSV-GFP based IFN-bioassay as we described before [3] and were diluted to a final concentration of 1000 UI/ml. The chIL28RA DF1 cell line, primed with recombinant chIFN-α or chIFN-λ or chIFN-γ or mock-treated with PBS for 24 hours, were subjected to RNA extraction. As an internal quality control, the transcription of mRNA for the chIFIT5 was quantified (data not shown) and samples that showed a significant induction of chIFIT5 were indicative of IFN-mediated stimulation of the gene [3]. The RNA sequencing was applied to investigate genes that are specifically or commonly induced by chicken type I, II and III IFNs (Supplementary Figure 1). A total of eight cDNA libraries, constructed from total RNA from IFN-stimulated chIL28RA DF1 cell, generated a total of 198,134,047 clean reads. These finished reads were mapped onto the chicken reference genome with an assembly rate of individual library ranged from 88.99% - 89.42%.

### 3.3. Comparative Characterization of ISGs Induced by Type I and Type III IFNs

Both type I (chIFN-α) and type III (chIFN-λ) IFNs bind to their specific receptors; however, they initiate the same downstream IFN-signalling pathways and culminate in the transcriptional activation of ISGs. The receptors for type I IFNs are ubiquitously expressed in majority of nucleated cells whereas the receptors for type III IFNs are more consolidated on epithelial cells. In order to investigate the nature of ISGs induced by specific types of IFN, we individually stimulated the established cells line and mapped the transcription of ISGs using transcriptomes.

Compared to the mock-treated chIL28RA DF1 cell, the chIFN-α stimulated a total of 111 genes and suppressed 29 genes whereas chIFN-λ up-regulated 115 and down regulated four genes (Figure 2A, Supplementary table). Using comparative fold induction criteria between chIFN-α and chIFN-λ, among genes that were differential expressed and mapped to the chicken genome, a total of 29 genes were observed to be chIFNα-specific whereas 60 were chIFN-λ-specific (Figure 2B). In order to understand the plasticity of chIFN-α and chIFN-λ in inducing specific set of ISGs, the comparative scatter plat analysis indicated that majority of the up-regulated as well as down-regulated genes were shared between type I and type III IFNs (Figure 2C) and thus represented a linear expression pattern. Next, we applied pairwise cluster plotting and noticed, as was observed in comparative dot plot, that genetics and nature of up- and down-regulated genes induced by chIFN-α and chIFN-λ were primarily aligned at both ends (Figure 2D). This distance matrix based pairing of IFN-regulated genes further confirms the linearity of the expression and highlighted the communal IFN-signalling pathways involved in response to type I and type III IFNs.

**Figure 2.**
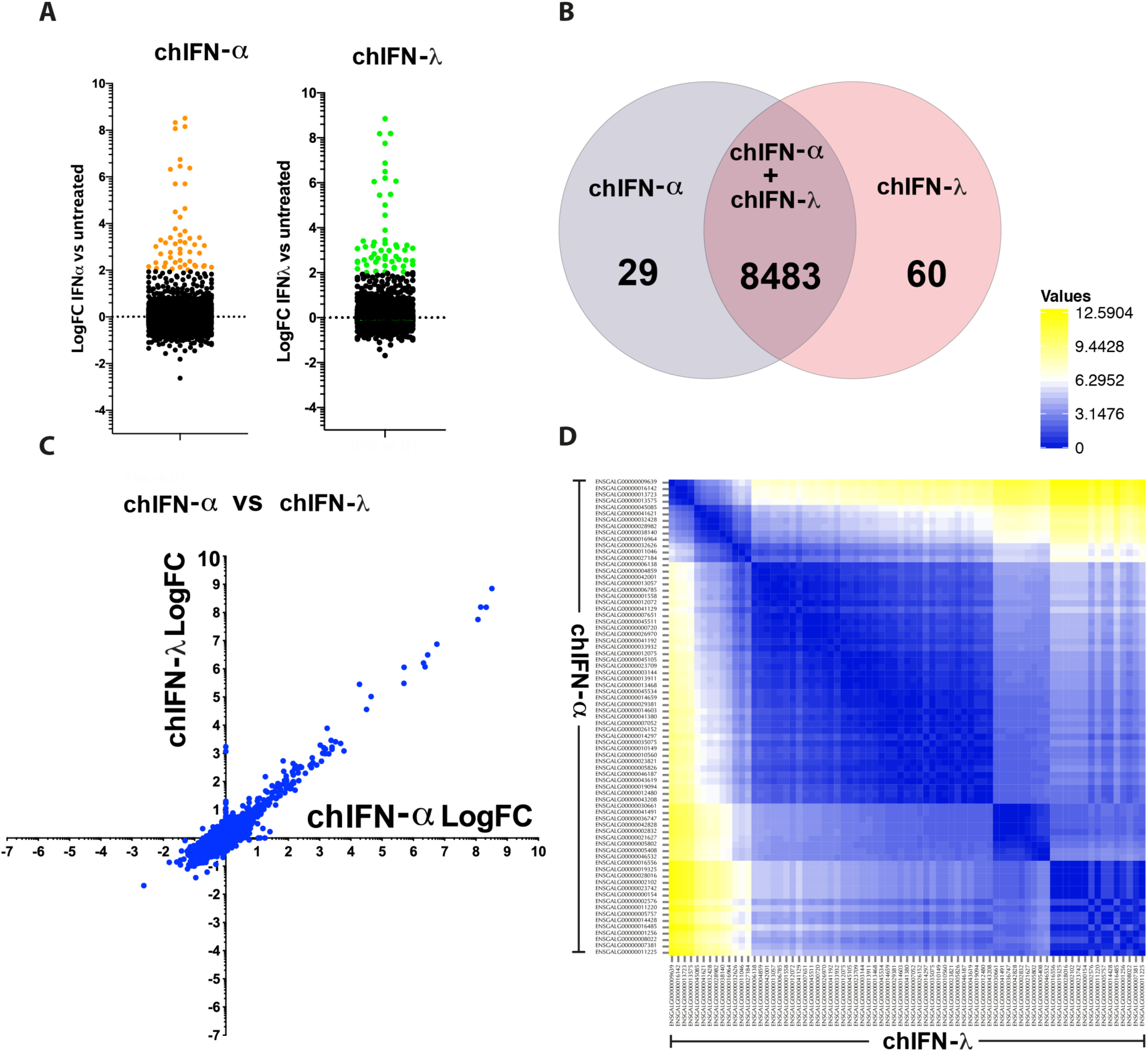
Expression patterns of type I and III IFN-regulated genes in chIL28RA DF1 cell line. (**A**) A number of ISGs were up-regulated and down-regulated by chIFN-α or chIFN-λ. Only significantly expressed genes with LogFC>2 compared to control are coloured green (**B**) Direct comparison of ISGs between both type I and type III IFNs revealed a regulation of common 8483 genes, and 29 and 60 specific to chIFN-α or chIFN-λ, respectively. (**C**) The dot plot representing the common and differential analysis of genes induced by chIFN-α or chIFN-λ. (**D**) Pairwise comparison of the genetic diversity of ISGs induced by type I and type III IFNs indicating a higher level of similarity between genes induced by chIFN-α or chIFN-λ.

### 3.4. Characteristic Induction of ISGs by Chicken Type I and Type II IFNs

Type I IFNs are encoded by multiple genes, however, there is only a single gene that translates into type II IFN (e.g. chIFN-γ). While type I IFN expression and functions are ubiquitous, the roles of type II IFNs are primarily restricted to the immune cells. Using a cell line that respond effectively to type I and type II IFNs, we mapped the nature and expression patterns of ISGs in chicken fibroblasts. Genes that were mapped to the chicken genome in chIFN-α and chIFN-γ-primed chIL28RA DF1 cell were 8449 and 8468, respectively (Supplementary table 1). Amongst these genes, a total of 110 were up-regulated and 36 were down-regulated by chIFN-γ in contrast to chIFNα-mediated up-regulation of 111 and down-regulation of 29 genes (Figure 3A, Supplementary table 1). The number of differentially regulated genes was comparable between chIFN-α and chIFN-γ-stimulated chicken fibroblasts, however, their nature and level of expression varied significantly. Direct comparison of genes between both type I and type II IFNs revealed a regulation of common 8448 genes, and 64 and 66 were specific to chIFN-α and chIFN-γ, respectively (Figure 3B).

**Figure 3.**
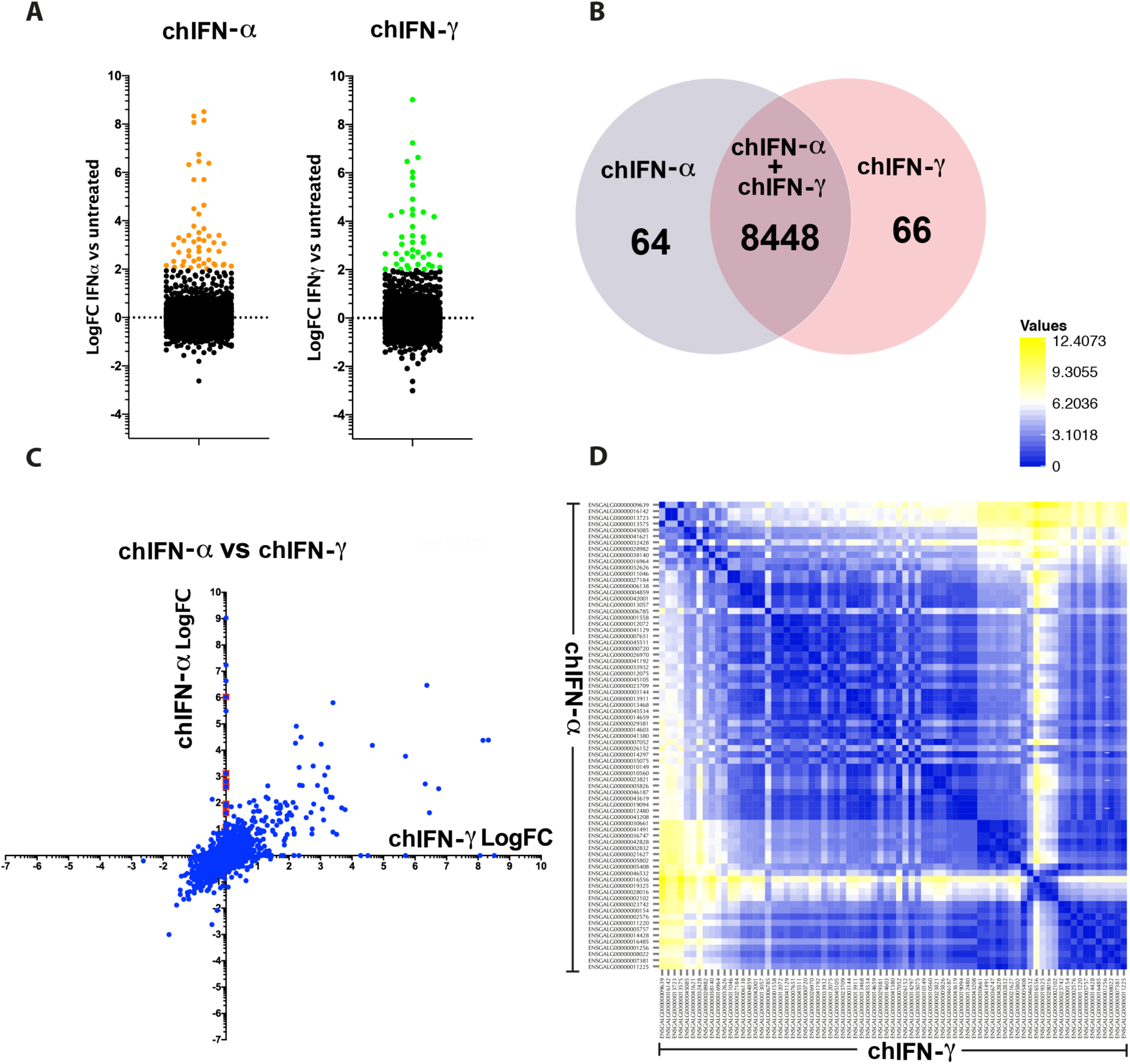
Diversity in the expression of type I and type II IFN-induced genes. (**A**) The dot plot shows the number of genes that we differentially regulated (up or down) by chIFN-α or chIFN-γ in chicken fibroblasts. Genes that are significantly up-regulated were marked green. (**B**) The Venn chart shows the number of genes induced by chIFN-α or chIFN-γ individually or mutually. (**C**) The scatter dot plot depicting the diversity in the expression of ISGs upon direct comparison between genes induced by chIFN-α or chIFN-γ. (**D**) Pairwise comparison of the significantly expressed genes between chIFN-α or chIFN-γ primed chicken fibroblasts. Genes with similar expression are coloured identical in the inter-section.

In order to map the diversity of these genes, we plotted them against the fold of induction of a gene by chIFN-α and compared it with the corresponding induction by chIFN-γ (Figure 3C). This direct comparison demonstrated the diversity patterns of ISGs. In contrast to comparison of ISGs induced by chIFN-α and chIFN-λ, a broader diversity of induction was observed between genes stimulated by chIFN-α and chIFN-γ. The pairwise comparison further displayed the genetic diversity of ISGs induced by type I and type II IFNs (Figure 3D). Altogether, these data demonstrate that differential interaction of type I and type II IFNs to their cognate receptors and induction of distinctive JAK-STAT mediated cascades led to a differential and characteristic pattern of ISGs by these IFNs.

### 3.5. Comparative Induction of ISGs by Type II and Type III IFNs

The receptors for the type II IFNs are predominantly expressed in majority of immune cells whereas receptors for type III IFNs are expressed in epithelial cells. While individual genes have been identified that are inducible by these cytokines, the global picture especially in cell lines to effectively respond to all analysed IFNs remained elusive. We observed a comparable number of genes up or down-regulated by chIFN-λ and chIFN-γ (Figure 4A). However, the genes were of diverse functionality and represented a range of signalling pathways, predominantly represented by ISGs (Figure 4A, Supplementary table). A total of 58 genes were significantly and differentially regulated by chIFNγ compared to 87 genes induced by chIFN-λ. Both IFNs, shared a great number of genes that were expressed and mapped to the annotated genome of the chicken (Figure 4B). While numbers of regulated genes were merely different, the nature and expression patterns were significantly variable. Majority of genes induced by chIFN-λ were also only weakly induced by chIFN-γ, however, a limited number of gene were equally induced by both chIFN-λ and chIFN-γ (Supplementary table 1, Figure 4C). Similar to the comparison of type I and type II, the pairwise scatting plot showed a range of distinctive expression patterns in ISGs that were induced by chIFN-λ and chIFN-γ (Figure 4D).

**Figure 4.**
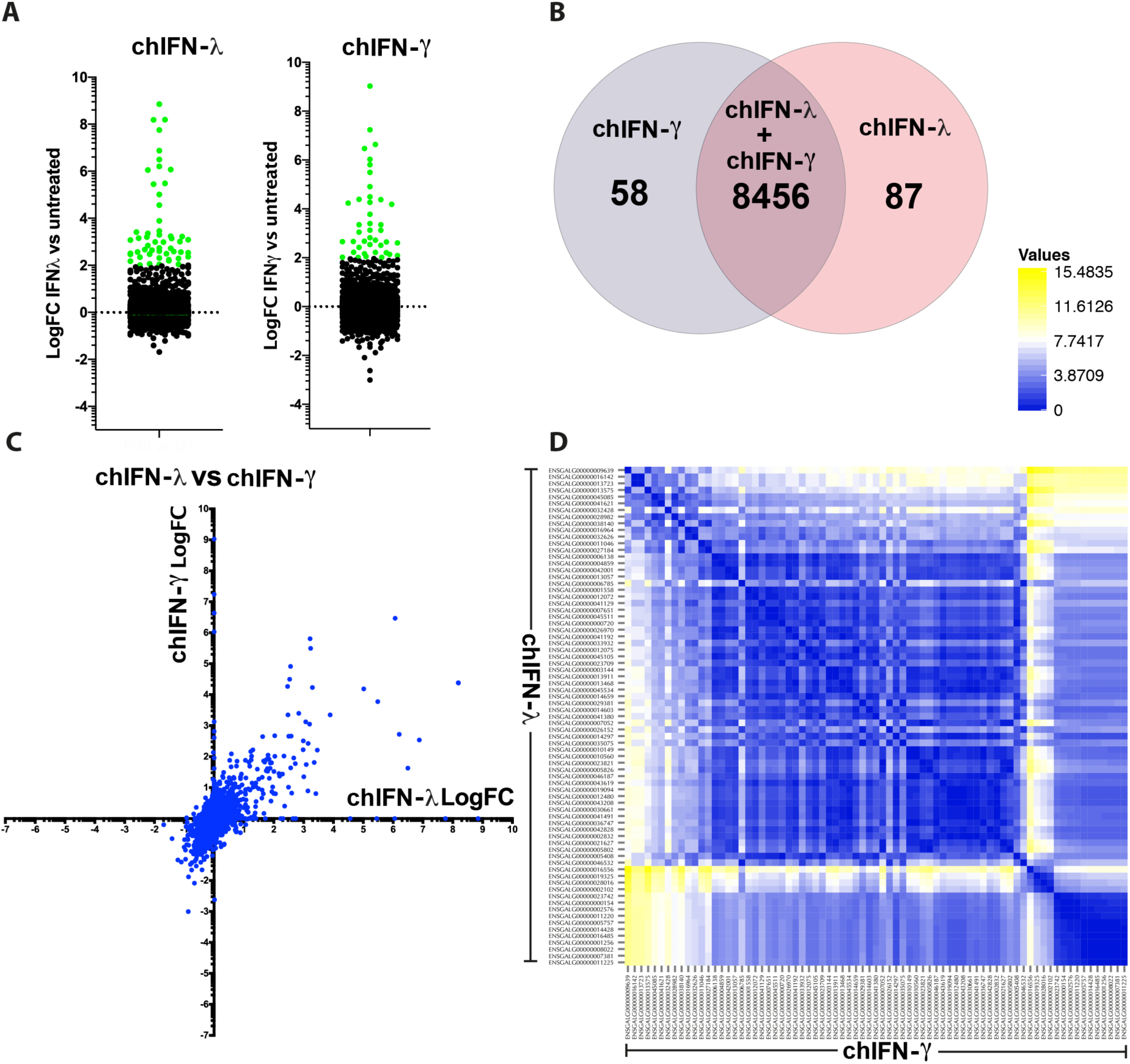
Comparative induction of ISGs by type II and type III IFNs. (**A**) Comparison of significantly (coloured green) up or down-regulation for ISGs by chIFN-λ and chIFN-γ. (**B**) A total of 58 ISGs were significantly and differentially regulated by chIFN-γ compared to 87 genes induced by chIFN-λ. Both IFNs shared a great number of genes that were expressed and mapped to the annotated genome of the chicken (n=8456). (**C**) Difference between the chIFN-λ and chIFN-γ induced genes revealing that majority of ISGs induced by chIFN-λ were weakly induced by chIFN-γ, however, there were a number of gene were equally induced by both chIFN-λ and chIFN-γ. (**D**) Pairwise scattering plot analysis showed a range of expression patterns in ISGs that were induced by chIFN-λ and chIFN-γ further indicating the difference in expression patterns.

### 3.6. Global and Differential Induction of ISGs by Chicken IFNs

Individual and combined analysis of genes induced by either type I or II or III IFNs concluded that each cytokine induced a range of genes in a cell line that express receptors for all types of IFNs (Figure 5A). However, the genetic and expression levels varied among genes that were induced by more than one IFN. While a large number of genes were insignificantly and ubiquitously expressed (n=8433), a range of commonalities and uniqueness in the expression dynamics was observed (Figure 5B). The Venn diagram of ISGs regulated by type I, II and III IFNs revealed majority of IFN-specific genes were expressed by the chIFN-γ followed by chIFN-λ and chIFN-α. However, as expected, the largest set of ISGs (n=50) was shared by the chIFN-λ and chIFN-α and the number of shared ISGs between type II and type III and type I and II were 23 and 15, respectively (Figure 5B).

**Figure 5.**
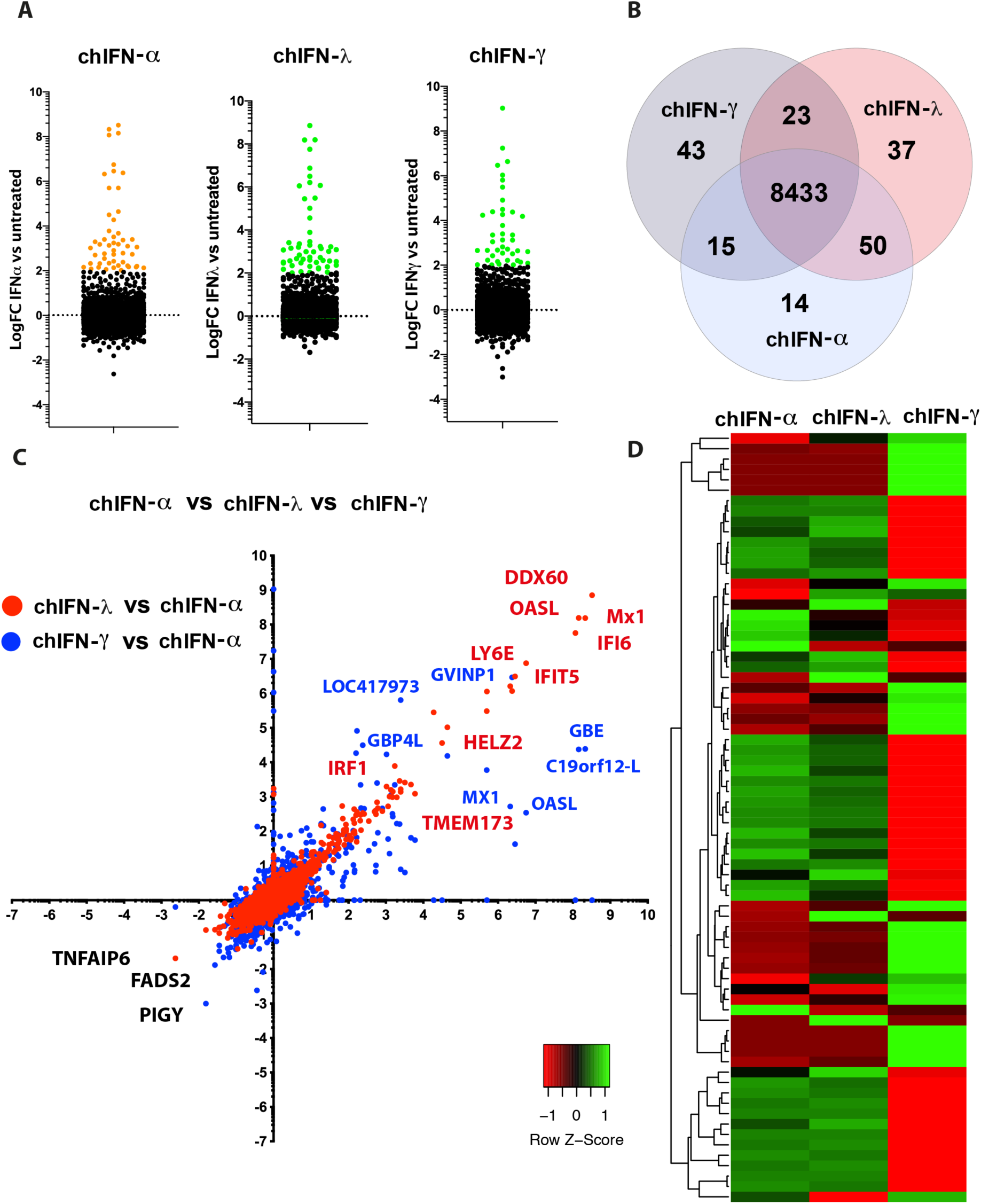
Genome-scale comparative analysis of ISGs induced by either type I or II or III. (**A**) Each IFN induced a range of genes in a cell line that express receptors for all types of IFNs. (**B**) The Venn diagram of ISGs regulated by type I, II and III IFNs revealed the largest set of ISGs (n=50) were shared by the chIFN-λ and chIFN-α while the number of shared ISGs between type II and type III and type I and II were 23 and 15, respectively. (**C**) Cumulative scatter plot analysis of ISGs between chIFN-λ and chIFN-α, and chIFN-γ and chIFN-α highlighted genes of overlapping and distinctive expression patterns. The prominent up-regulated and down-regulated are labelled and colour-coded. (**D**) Heat map to compare the selected un-regulated ISGs induced by chIFN-α, chIFN-γ and chIFN-λ indicates less-extensive expression divergence between genes of type I and type III IFNs, whereas the nature of type II IFN-mediated genes showed a higher divergence with the genes induced by the rest of IFNs.

Cumulative scatter plot analysis of genes between chIFN-λ and chIFN-α, and chIFN-γ and chIFN-α highlighted genes of overlapping and distinctive expression patterns. The genes that were up-regulated ranged from transcription factors, nuclear receptors and host antiviral factors. However, these genes were predominantly presented by the ISG including DDX60, OASL, Mx1, IFI6, IFIT5, LY6E, TMEM173 among others (Figure 5C). While the down-regulated genes were lower in numbers, transcription of genes such as PIGY, FADS and TNFAIP6 were unanimously suppressed by IFNs in chicken fibroblasts (Supplementary table 2). The clustering heat map, directly comparing selecting unregulated genes among chIFN-α, chIFN-γ and chIFN-λ, indicated a distribution of a set of genes across the spectrum. This heat map articulated a less-extensive expression divergence between genes of type I and type III IFNs, whereas the nature of type II IFN-mediated genes showed a higher divergence with the genes induced by the rest of IFNs (Figure 5D). Altogether, this data highlight commonalities and fundamental differences in the potential signalling cascades that drive the expression of ISGs.

### 3.7. Promoter in trans Analysis of Differentially Regulated ISGs

The direct comparative analysis of significantly regulated genes showed a high-level of synergy between chIFN-α and chIFN-γ in both up- and down-regulated genes where PIGY was the most down-regulated and DDX60 was the most up-regulated gene (Figure 6A). However, both up- and down-regulated genes were substantially different between type I and II, and type II and type III IFNs-stimulated genes. This differential and IFN-dependent expression of ISGs is mainly attributed to the promoter of the genes [2,3]. Next, we scanned the chicken genome for the presence of ISRE and GAS elements in top regulated genes and identified that highly regulated genes carried GAS element in abundance compared to the ISRE in an IFN-independent manner (Table 1, Supplementary table 3). The sequence of ISRE and GAS varied among genes induced by IFNs where GAS element was identified in type I and type III ISGs and even the ISRE was detected in the promoter of genes induced by type II IFNs with a consensus sequence shown in the Figure 6B and 6C for ISRE and GAS, respectively.

**Table 1.**
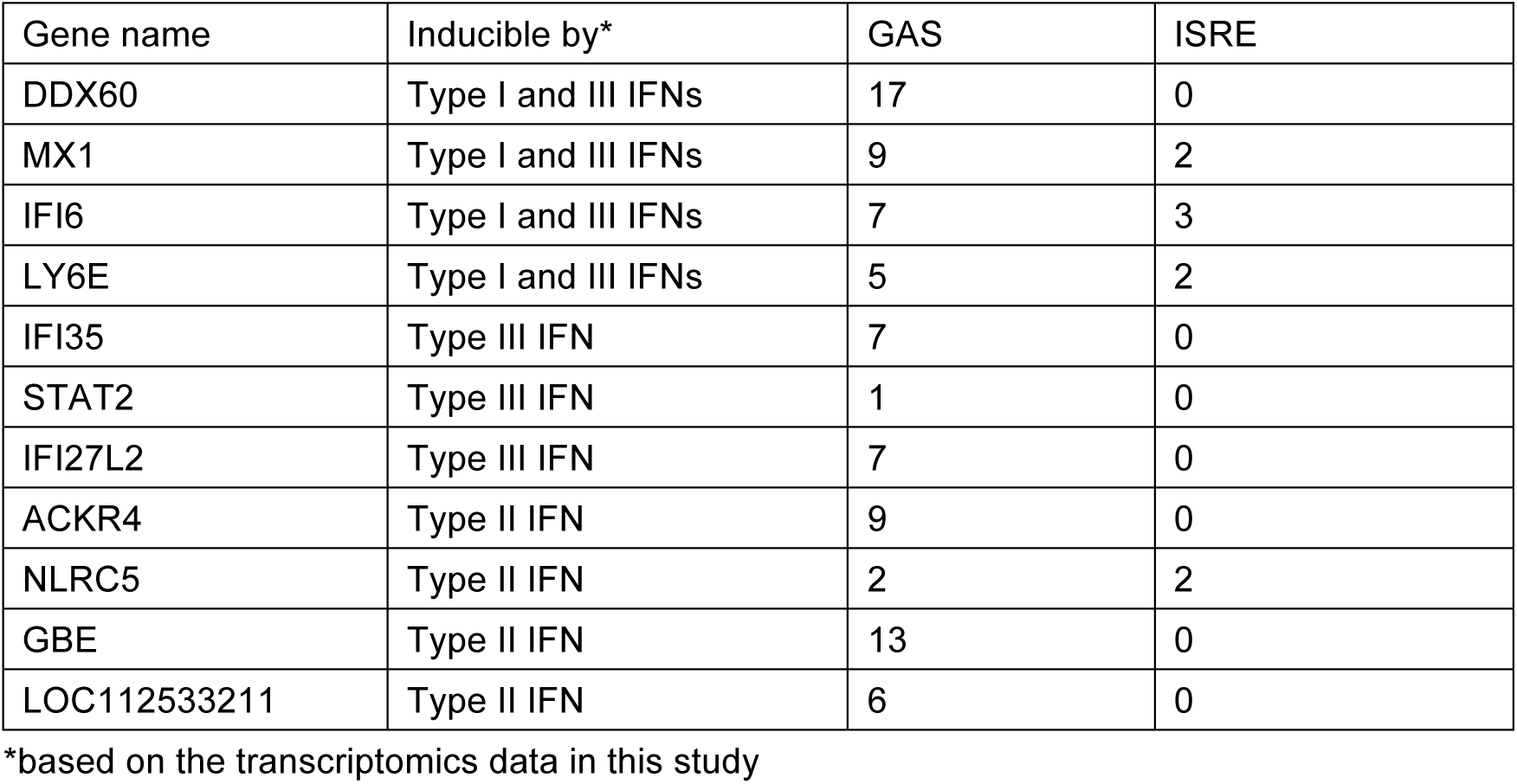
Number of ISRE and/or GAS in selected ISGs.

**Figure 6.**
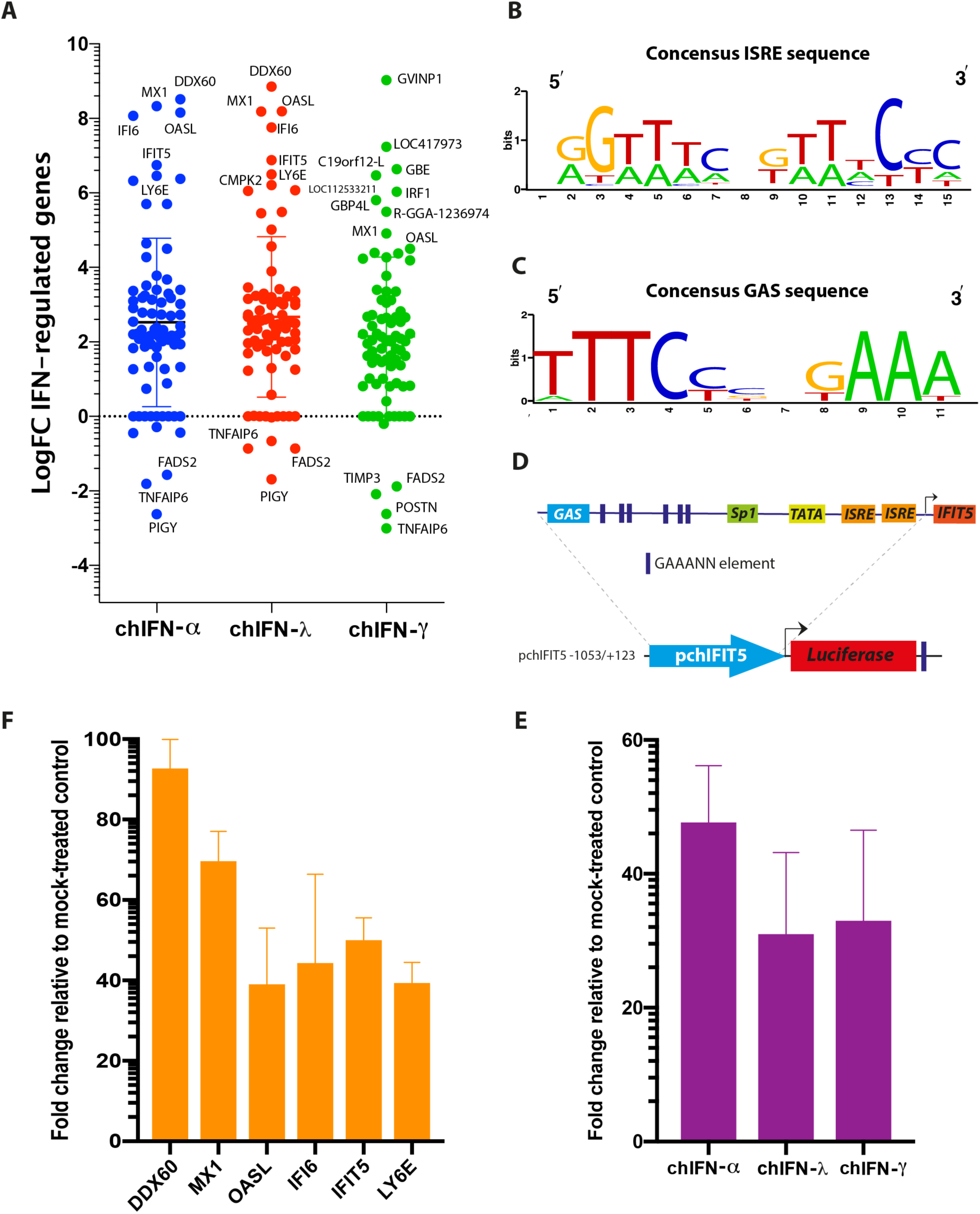
Expression of genes individually regulated by type I, II and III IFNs and their association with GAS and ISRE elements in the promoter region. (**A**) Direct comparative analysis of significantly regulated ISGs showed a high-level synergy between chIFN-α and chIFN-γ in both up- and down-regulated genes; DDX60 was the most up-regulated gene while PIGY was the most down-regulated. ISRE (**B**) and GAS (C) consensus sequence in the most up-regulated ISGs. (**D**) The promoter sequence of chIFITI5 (pchIFIT5) contained two putative ISREs, a GAS and six GAAANN elements. (**E**) The chIL28RA DF-1 cells were transfected with pchIFIT5 and stimulated with chIFN-α, chIFN-γ or chIFN-λ. The fold change induction in the luciferase genes compared to mock-primed cells is plotted. A substantial stimulation of reporter gene transcription by all type I, II and III IFNs was observed. (**F**) Fold change induction of significantly up-regulated ISGs using qPCR.

In order to demonstrate that *in silico* predicted ISRE and GAS respond to IFNs, we generated a luciferase reporter system using wild type promoter of chIFIT5, which is one of the highly regulated genes both transcriptionally and translationally in chicken [3]. The promoter sequence of the chIFITI5 contained two putative ISREs, a GAS and six GAAANN elements, which are hallmarks motifs for IFN-responsiveness (Figure 6D). This entire cassette was cloned upstream to luciferase reported gene and was used to monitor the expression of the gene in cells primed individually with chIFN-α, chIFN-λ or chIFN-γ. Fold change induction, compared to mock-primed cells, showed a substantial stimulation of reporter gene transcription by all type I, III and II IFNs (Figure 6E). Finally, we verified the RNA-seq based expression of top six ISGs using qPCR as we described before [3], and the fold change of all these ISGs aligned with the level of induction observed in transcriptomes (Figure 6F).

## 4. Discussion

Interferons were first described in “chicken” when Isaacs and Lindenmann noticed that supernatant from virus-infected cells can “interfere” with the replication of influenza virus [29]. However, vast majority of information on the nature of IFNs regulated and IFN-pathways in chicken is derived from studies conducted in human and mice. Moreover, substantial differences in several innate immune checkpoints have been observed in chickens, which highlight fundamental and evolutionary distinct mechanisms in birds especially in chickens [3]. Thus, understanding mechanisms and gaining insights into additional compensatory mechanisms in IFN-mediated establishment of antiviral state in chicken would provide foundations to underpin diversity and dynamics of innate immunity in animals.

Since ISGs play the most dynamic and versatile roles in inhibiting viruses at multiple scales, genetic annotation of these genes is essential to perform large-scale functional studies and to reveals functional plasticity of ISGs. While information in mammals, especially in human and mice, has started to expand in unraveling the roles of ISGs, studies have proposed the expression of ISGs both in vitro [30] and in vivo [18] in chicken; however, this information is only limited to type I IFNs or to a limited set of type III IFNs [31]. At system and cellular levels, IFNs and their regulated genes function in a network and thus necessitate the exploration of these components at the genome-wide scale. Moreover, mapping and cataloguing genes that are specific to type I, II or III IFNs or identifying genes that span the responses against pan-IFN levels would provide foundation to functional investigations against viruses of birds.

Building upon existing information on chicken ISGs [18,30,32] and our recent studies [3, 33], we compared the expression of the genes, which are induced by either chIFN-α, chIFN-λ or chIFN-γ and catalogue the genes which are IFN-specific or genes that are shared between different types of IFNs in chicken. Since IFNs show cell and tissue-specific actions [8], which are mainly attributed to their cognate expression patterns [28], we attempted to establish a model system that could reliably respond to all type of IFNs. The chicken fibroblast (DF-1), one of the most characterized cell lines in avian research, was selected. However, due to repressive expression of chIFN-λ receptors [28], the treatment of DF1 with type III IFN showed defective genes inductions, IFN-promoter activation and cumulatively established an antiviral state against IFN-sensitive virus (VSV). We thus exploited a permanent DF1 cell line that exogenously express chIFN-λ receptors and thus revised the full responsiveness to type III as well as other IFNs. We propose this as an improved tool to study genome-scale ISGs induced by multiple stimuli and regulated by a complex cellular network.

Using genome-scale transcriptomes in chicken fibroblasts, we enlisted genes that were IFN-specific or were generic. The overall numbers of genes that were either up- or down-regulated were lower compared to previous studies which were performed on primary [17] or established cell lines or *in vivo* [18]. We reasoned that since we applied stringent-selection and fold-induction criteria in a cell line that could induce simultaneous responses, the overall gene transcription was weaker. Moreover, the *in vitro* experimental setting would differ from a naturally occurring infection where more potent local type I IFNs are activated soon after viral infections (or other stimuli) which then expand to system with the release of several cytokines and other affecters simultaneously [34]. However, significantly up- or down-regulated genes in our study were known IFN-stimulated (e.g. ISGs). Moreover, chIFIT5, which we have previously identified and characterized [3], showed higher expression in type I and type III-treated chicken fibroblasts. Additionally, owing to the fact that different cells are known to respond IFNs differently [4] and that ISGs express temporally depending on the nature of stimuli [18], gauging the expression dynamics at different time points with both viral and IFNs stimulated cell would provide complete picture in future.

The direct comparison of all ISGs regulated by type I, II and III in heatmap and scatter plots indicated that the effects of chIFN-α and chIFN-λ on the expression of ISGs were distinctive from the genes expressed in chIFN-γ-stimulated cells. As anticipated, a higher resolution analysis indicated that the differentially expressed genes included a list of known genes such as IFIT5, Mx1, OASL further supporting the reliability of experimental approach and suitability of the model. However, as Roll et al. 2018 [18] have highlighted it, yet poor annotation of the chicken genome may lead to either false gene annotation or lack of gene assembly results in missed genes in the list of otherwise IFN-stimulated genes.

During the analysis of IFN-mediated gene expression, it was revealed that up-regulated genes were predominantly presented over the down-regulated genes in all tested type I, II and III IFNs. It was only PIGY (phosphatidylinositol glycan anchor biosynthesis) that was significantly down regulated by the type I IFN and three genes named TNFAIP6 (Tumor Necrosis Factor-Inducible Gene 6 Protein), POSTN (Periostin) and TIMP3 (TIMP Metallopeptidase Inhibitor 3) were down regulated by the type II IFNs. One plausible rationale for higher number of upregulated genes is the trait of all IFNs and is supported by the presence of ISRE and GAS elements in majority of analysed ISGs. Analysis of highly expressed ISGs in different IFN-treated lists revealed presence of a number of ISRE and GAS motifs. Intriguingly and in contrast to human promoter sequences [27], a large number of GAS sequences were identified compared to ISRE motifs in chicken ISGs. Comparing these motifs with the nature of treated IFNs, no correlation was observed; all types of IFN-regulated genes carried both ISRE and GAS elements. While we demonstrated that chicken IFNs transcriptionally induced a reporter gene through GAS and ISRE in an IFN-independent manner, it requires future experimental studies to identify the biological relevance of *in silico* predicted motifs with the functional relevance. The transcriptomes and individual reports on ISGs could not clearly distinguish between primary (direct IFN-dependents) and secondary (positive loop feedback of IFNs) responses, thus impeding a clear differentiation of genes expressions that are specific to each of these stimuli. Moreover, majority of functional studies have focused on the up-regulated genes, the potential anti- or pro-viral roles of under-expressed genes would shed lights on the dynamics and plasticity of IFN regulated genes.

Owing to the fact that ISGs play fundamental roles in a wide range of cellular activities including transcriptional and translational regulation of immune responses [35,35] and establishing a host antiviral state against viruses, the provided data would lay foundations to investigate functions of these important and complex domains of host-pathogen interactions using large-scale screening platforms. This comprehensive information is first of its kind to compare genes that are differentially regulated by type I, II and type III IFNs in chicken.

## Acknowledgments

We appreciate the kind gift of pchMx-luc by Nicolas Ruggli, Switzerland, and VSV-GFP and chIL28RA DF-1 by Dennis Rubbenstroth, Institut für Virologie, Universitätsklinikum Freiburg, Germany.

## Data Availability

The data generated in this study were submitted in the Bioproject database under ID: PRJNA678234 and PRJNA678244.

## References

1. Pestka, S.; Krause, C.D.; Walter, M.R. Interferons, interferon-like cytokines, and their receptors. Immunol. Rev. 2004, 202, 8–32.

2. Santhakumar, D.; Rubbenstroth, D.; Martinez-Sobrido, L.; Munir, M. Avian Interferons and Their Antiviral Effectors. Front. Immunol. 2017a, 8:49.

3. Santhakumar, D.; Rohaim, M.A.S.; Hussein, H.A.; Hawes, P.; Ferreira, H.L.; Behboudi, S.; Iqbal, M.; Nair, V.; Arns, C.W.; Munir, M. Chicken Interferon-induced Protein with Tetratricopeptide Repeats 5 Antagonizes Replication of RNA Viruses. Sci. Rep. 2018, 8(1): 6794.

4. Santhakumar, D.; Iqbal, M.; Nair, V.; Munir, M. Chicken IFN Kappa: A Novel Cytokine with Antiviral Activities. Sci. Rep. 2017b, 7(1): 2719.

5. Fensterl, V.; Sen, G.C. Interferons and viral infections. Biofactors 2009, 35(1):14–20.

6. Lazear, H.M.; Schoggins, J.W.; Diamond, M.S. Shared and Distinct Functions of Type I and Type III Interferons. Immunity 2019, 50(4):907–923.

7. Randall, R.E.; Goodbourn, S. Interferons and viruses: an interplay between induction, signalling, antiviral responses and virus countermeasures. J. Gen. Virol. 2008, 89, 1–47.

8. Kessler, D.S., Veals, S.A., Fu, X.Y.; Levy, D.E. Interferon-alpha regulates nuclear translocation and DNA-binding affinity of ISGF3, a multimeric transcriptional activator. Genes. Dev. 1990, 4(10): 1753–65.

9. Schoggins, J.W. Recent advances in antiviral interferon stimulated gene biology. F1000Res. 2018, 7:309.

10. Schoggins, J.W.; Rice, C.M. Interferon-stimulated genes and their antiviral effector functions. Curr. Opin. Virol. 2011a, 1(6):519–25.

11. Schoggins, J.W.; Wilson, S.J.; Panis, M.; Murphy, M.Y.; Jones, C.T.; Bieniasz, P.; Rice, C.M. A diverse range of gene products are effectors of the type I interferon antiviral response. Nature. 2011b, 472(7344): 481–5.

12. Ramos, I.; Smith, G.; Ruf-Zamojski, F.; Martínez-Romero, C.; Fribourg, M.; Carbajal, E.A.; Hartmann, B.M.; Nair, V.D.; Marjanovic, N.; et al. Innate Immune Response to Influenza Virus at Single-Cell Resolution in Human Epithelial Cells Revealed Paracrine Induction of Interferon Lambda. J. Virol. 2019, 90: 20, e00559–19.

13. USDA. Available from: https://www.aphis.usda.gov/aphis/ourfocus/animalhealth/animal-disease-information/avian-influenza-disease/ct_avian_influenza_disease

14. Uchikawa, E.; Lethier, M.; Malet, H.; Brunel, J.; Gerlier, D.; Cusack, S. Structural analysis of dsRNA binding to anti-viral pattern recognition receptors LGP2 and MDA5. Mol. Cell. 2016, 62(4):586–602.

15. Karpala, A.J.; Bingham, J.; Schat, K.A.; Chen, L.M.; Donis, R.O.; Lowenthal, J.W.; Bean, A.G. Highly pathogenic (H5N1) avian influenza induces an inflammatory T helper type 1 cytokine response in the chicken. J. Interferon. Cytokine. Res. 2011, 31(4):393–400.

16. Liniger, M.; Summerfield, A.; Zimmer, G.; McCullough, K.C.; Ruggli, N. Chicken cells sense influenza A virus infection through MDA5 and CARDIF signaling involving LGP2. J. Virol. 2012, 86(2):705–17.

17. Giotis, E.S.; Robey, R.C.; Skinner, N.G.; Tomlinson, C.D.; Goodbourn, S.; Skinner, M.A. Chicken interferome: avian interferon-stimulated genes identified by microarray and RNA-seq of primary chick embryo fibroblasts treated with a chicken type I interferon (IFN-α). Vet. Res. 2016, 47(1):75.

18. Roll, S.; Hartle, S.; Lutteke, T.; Kaspers, B.; Hartle, S. Tissue and time specific expression pattern of interferon regulated genes in the chicken. BMC Genomics 2017, 18, 264.

19. Berger Rentsch, M.; Zimmer, G. A vesicular stomatitis virus replicon-based bioassay for the rapid and sensitive determination of multi-species type I interferon. PLoS One. 2011, 6, e25858.

20. Bolger, A.M.; Lohse, M.; Usadel, B. Trimmomatic: a flexible trimmer for Illumina sequence data. Bioinformatics. 2014, 30(15): 2114–20.

21. Dobin, A.; Davis, C.A.; Schlesinger, F.; Drenkow, J.; Zaleski, C.; Jha, S.; Batut, P.; Chaisson, M.; Gingeras, T. R. STAR: ultrafast universal RNA-seq aligner. Bioinformatics, 2013, 29(1): 15–21.

22. Robinson, M.D.; McCarthy, D.J.; Smyth, G.K. edgeR: a Bioconductor package for differential expression analysis of digital gene expression data. Bioinformatics. 2010, 26(1): 139–140.

23. Li, W.; Freudenberg, J.; Suh, Y.J.; Yang, Y. Using volcano plots and regularized-chi statistics in genetic association studies. Computational Biology and Chemistry. 2014, 48: 77–83.

24. Subhash, S.; Kanduri, C. GeneSCF: a real-time based functional enrichment tool with support for multiple organisms. BMC Bioinformatics. 2016, 17(1):365.

25. Anders, S.; Reyes, A.; Huber, W. Detecting differential usage of exons from RNA-seq data. Genome Res. 2012, 22(10): 2008–17.

26. Schumacher, B.; Bernasconi, D.; Schultz, U.; Staeheli, P. The chicken Mx promoter contains an ISRE motif and confers interferon inducibility to a reporter gene in chick and monkey cells. Virology. 1994, 203(1):144–8.

27. Tsukahara, T.; Kim, S.; Taylor, M.W. REFINEMENT: a search framework for the identification of interferon-responsive elements in DNA sequences--a case study with ISRE and GAS. Comput. Biol. Chem. 2006, 30(2):134–47.

28. Reuter, A.; Soubies, S.; Härtle, S.; Schusser, B.; Kaspers, B.; Staeheli, P.; Rubbenstroth, D. Antiviral activity of lambda interferon in chickens. J. Virol. 2014, 88(5):2835–43.

29. Isaacs, A.; Lindenmann, J. Virus interference. I. The interferon. Proc. R. Soc. Lond. Ser. B Biol. Sci. 1957, 147, 258–267.

30. Giotis, E.S.; Ross, C.S.; Robey, R.C.; Nohturfft, A.; Goodbourn, S.; Skinner, M.A. Constitutively elevated levels of SOCS1 suppress innate responses in DF-1 immortalised chicken fibroblast cells. Sci. Rep. 2017, 7(1):17485.

31. Masuda, Y.; Matsuda, A.; Usui, T.; Sugai, T.; Asano, A.; Yamano, Y. Biological effects of chicken type III interferon on expression of interferon-stimulated genes in chickens: comparison with type I and type II interferons. J. Vet. Med. Sci. 2012, 74, 1381–1386.

32. Xie, T.; Chen, T.; Li, C.; Wang, W.; Cao, L.; Rao, H.; Yang, Q.; Shu, H.B.; Xu, L.G. RACK1 attenuates RLR antiviral signaling by targeting VISA-TRAF complexes. Biochem Biophys Res Commun. 2019, 508(3):667–674.

33. Arslan, M.; Yang, X.; Santhakumar, D.; Liu, X.; Hu, X.; Munir, M.; Li, Y.; Zhang, Z. Dynamic Expression of Interferon Lambda Regulated Genes in Primary Fibroblasts and Immune Organs of the Chicken. Genes (Basel). 2019, 10(2). pii: E145.

34. Penski, N.; Penski, N.; Härtle, S.; Rubbenstroth, D.; Krohmann, C.; Ruggli, N.; Schusser, B.; Pfann, M.; Reuter, A.; Gohrbandt, S.; et al. Highly pathogenic avian influenza viruses do not inhibit interferon synthesis in infected chickens but can override the interferon-induced antiviral state. J. Virol. 2011, 85(15):7730–41.

35. de Veer, M.J.; Holko, M.; Frevel, M.; Walker, E.; Der, S.; Paranjape, J.M.; Silverman, R.H.; Williams, B.R. Functional classification of interferon-stimulated genes identified using microarrays. J. Leukoc. Biol. 2001, 69 (6):912–20.

36. To, K.K.; Lau, C.C.; Woo, P.C.; Lau, S.K.; Chan, J.F.; Chan, K.H.; Zhang, A.J,; Chen, H.; Yuen, K.Y. Human H7N9 virus induces a more pronounced pro-inflammatory cytokine but an attenuated interferon response in human bronchial epithelial cells when compared with an epidemiologically-linked chicken H7N9 virus. Virol. J. 2016, 13:42 (2016)

